# A library of cell-surface and secreted proteins from *Schistosoma mansoni* identifies early serological markers of infection

**DOI:** 10.1101/593616

**Authors:** Cécile Crosnier, Anna V. Protasio, Gabriel Rinaldi, Shona Wilson, Matthew Berriman, Gavin J. Wright

## Abstract

Schistosomiasis is a major global health problem caused by blood-dwelling parasitic worms and is currently treated by the mass administration of the drug praziquantel. Appropriate drug treatment strategies are informed by diagnostics that establish the prevalence and intensity of infection, which, in regions of low transmission should be highly sensitive. To identify sensitive new serological markers of *Schistosoma mansoni* infections, we have compiled a recombinant protein library of 115 parasite cell surface and secreted proteins expressed in mammalian cells. The vast majority of them were shown to be immunoreactive and to contain heat-labile conformational epitopes when tested against pooled human sera from endemic regions. After probing the library against a time series of sera samples from experimental infections in mice, we identified several markers of infection, the majority of which belong to the saposin-domain-containing and cathepsin families of proteins. These new markers will be a valuable tool to detect ongoing and previous *S. mansoni* infections, including in regions of low transmission. We envisage that this new recombinant protein resource will be used in a wide range of cellular and molecular assays to further our understanding of *Schistosoma* biology.

## Introduction

Schistosomiasis is a parasitic disease affecting over 200 million people in 52 countries and is considered one of the world’s major health problems causing 200,000 deaths per year. In 2015, the impact of the disease was estimated at 3.5 million disability-adjusted life years (DALYS), putting a huge socio-economic burden on many low and middle-income countries [1]. In humans, schistosomiasis is caused by five species of platyhelminth parasite belonging to the genus *Schistosoma*. Their geographical distribution is restricted by the presence of species-specific freshwater snails that act as intermediate hosts, with *S. mansoni* having the most widespread distribution, encompassing both Africa and South America [2]. Infected snails shed cercariae, a free-swimming form of the parasite, which penetrates human skin to initiate infection. Maturation of the parasite within its human host takes several weeks: cercariae first remodel their surface to form schistosomula [3, 4], which migrate through the dermis for several days before entering the host bloodstream. Eventually, young adult male and female *S. mansoni* worms pair up in the liver before moving to the mesenteric blood vessels, where each pair can release over 300 eggs per day from around five weeks post-infection [5]. The symptoms of the disease are caused by progressive accumulation of eggs within host tissues which elicit host-derived inflammatory responses that can eventually lead to liver fibrosis, portal hypertension and, if left untreated, death [6]. In the absence of a licenced vaccine, preventative treatment relies on the use of a single drug, praziquantel.

The exact prevalence of schistosomiasis worldwide may be underestimated because of the limitations of routine methods of detection [7]. Diagnosis of *S. mansoni* infections and subsequent decisions on mass drug administration mainly rely on observing parasite eggs in the stools of patients: the Kato-Katz test. Although this method can detect current infections, it is not sufficiently sensitive to diagnose low levels of infection present in areas of low-endemicity, or in recently-treated populations [8, 9]. In these instances, the direct detection of parasite antigens such as the circulating anodic antigen (CAA) in patients’ sera is a more sensitive method [10]. In areas where elimination has been achieved, the detection of anti-parasite antibodies in patients’ sera is a good approach to monitor historical exposure to the parasite, thus avoiding any risk of resurgence. Host antibody responses to *S. mansoni* parasites is often measured against whole parasite extracts such as soluble egg antigens (SEA) or soluble worm antigen preparation (SWAP), which are not molecularly defined and can lead to cross-reactivity with other helminths species. While a diagnostic test based on a recombinant protein could mitigate these problems, only very few have been used to test patient antibody responses to *Schistosoma* infections [11]. The development of new, more sensitive diagnostic tools would therefore help to improve the detection and early treatment of schistosomiasis.

Extracellular antigens released by or displayed at the surface of the parasite at the initial stage of infection can be valuable early immunodiagnostic markers as they are directly exposed to the host humoral system [3]. The identification of such antigens has been aided by the sequencing and annotation of the *S. mansoni* genome [12] and several proteomics [13–21], transcriptional [22–27] and *in silico* analyses [28], which have identified genes expressed by the schistosomula and adult worm. Despite their value, extracellular proteins pose challenges for recombinant expression because they contain structurally important posttranslational modifications such as glycosylation and especially disulphide bonds, which are often required to produce informative antibody epitopes. To address this, we have previously developed protein expression approaches using a mammalian system that can be used to compile large panels of parasite secreted recombinant ectodomains that retain their binding activity and immunogenicity [29, 30], enabling the identification of host-parasite receptor-ligand interactions and humoral markers of protection against malaria [31].

To identify new markers for *S. mansoni* infections, we created a panel of 115 recombinant proteins representing secreted and membrane-tethered *S. mansoni* proteins that were mostly enriched for expression at the schistosomula stage. Using mouse sera from experimentally-controlled *S. mansoni* infections, we were able to identify several early serological markers of infection mostly belonging to the saposin-containing and cathepsin families of protein.

## Material and methods

### Identification of S. mansoni cell-surface and secreted proteins

Genes encoding cell surface and secreted proteins from *S. mansoni* were identified using published proteomic and transcriptional data [13–16, 19–23]. To enrich for genes transcribed at the schistosomula stage, we identified 1,302 transcripts upregulated at 48 compared to 3 hours posttransformation [32] using EdgeR [33] as well as the 1,000 most abundant transcripts at 48 hours as assessed by reads per kilobase mapped, resulting in 1,977 unique genes. Genes encoding 274 secreted and cell surface proteins were identified from this list using signal peptide and transmembrane domain prediction software [34, 35]. Proteins spanning the membrane multiple times and therefore difficult to express as a contiguous ectodomain, and those with sequence homologies to mitochondrial or endoplasmic reticulum proteins were subsequently excluded, leaving a shortlist of 180 proteins. Gene structures were manually refined by mapping transcriptome data to the genome sequence, and those genes which spanned gaps in the genome sequence, or with ambiguity in the gene structure (e.g. due to genome misassemblies) were removed, resulting in a final list of 115 candidates. Because most proteins were previously uncharacterised and therefore known only by their database accession numbers, all proteins described in this study were given numbers, which we use throughout the text for convenience (Table 1).

**Table 1.** Details of 115 cell-surface and secreted proteins from *Schistosoma mansoni*. For each protein, are provided from left to right the accession number, alternative name, boundaries of the extracellular domain expressed in mammalian cells, expected molecular mass in kiloDaltons (MM), domain similarities and level of expression as determined by ELISA. References to previous proteomics studies where the proteins were identified are provided where available. The expected molecular mass of each protein was calculated by adding 3kDa per predicted glycosylation site (the average mass of a N-linked glycan) to the expected mass of the protein. Levels of expression were determined from each ELISA profile as detailed in the Material and Methods section. IgSF, immunoglobulin superfamily protein.

| Number | Accession # | Name | Boundaries | MM | Domain/Protein similarity | Level | Ref. |
| --- | --- | --- | --- | --- | --- | --- | --- |
| Cell surface |  |  |  |  |  |  |  |
| 1 | Smp_195190 | Sm13 | E18-T80 | 37 |  | High | [17] |
| 2 | Smp_081920 | SmLy6I (Cd59.5) | L26-T104 | 45 |  | Medium | [17] |
| 3 | Smp_166340 | SmLy6F (Cd59.4) | L26-S98 | 47 |  | Medium | [17] |
| 4 | Smp_017730 | Sm200 | D20-S1662 | 257 |  | Medium | [14] |
| 5 | Smp_127820 |  | V18-S760 | 132 |  | Medium | [14] |
| 6 | Smp_194920 |  | D17-T592 | 98 | T cell immunomodulatory protein | Low | [14] |
| 7 | Smp_011680 |  | L30-P348 | 87 | Cd36-like class B scavenger receptor | Low |  |
| 8 | Smp_054070 |  | D31-S210 | 71 | TM2 domain-containing protein 3 | High |  |
| 9 | Smp_073400 |  | R25-T214 | 78 | LAMP-like protein | Medium |  |
| 10 | Smp_105220 | SmLyB (Cd59.2) | I20-P99 | 42 |  | High | [15] |
| 11 | Smp_019350 | SmLy6A (Cd59.1) | H28-T102 | 38 |  | High | [15] |
| 12 | Smp_021220 |  | E24-S119 | 53 |  | High |  |
| 13 | Smp_031880 |  | R18-S240 | 58 | Ig domain-containing protein, basigin-related | High |  |
| 14 | Smp_009830 |  | D20-S149 | 51 | Translocon-associated protein subunit beta | Low |  |
| 15 | Smp_032520 |  | Y19-R200 | 74 | LAMP-like protein | High |  |
| 16 | Smp_074000 |  | I19-I232 | 56 |  | Low |  |
| 17 | Smp_102480 |  | N29-D66 | 37 |  | High |  |
| 18 | Smp_124000 | MEG-14 | T27-D110 | 38 |  | High |  |
| 19 | Smp_156270 |  | S23-P144 | 47 | Post-GPI attachment to protein factor | Low |  |
| 20 | Smp_174580 |  | S17-T275 | 59 | Vesicular integral membrane protein | High |  |
| 21 | Smp_176020 | MEG-11 | D23-P55 | 34 |  | High |  |
| 22 | Smp_048380 |  | L20-S283 | 73 |  | Medium |  |
| 23 | Smp_060570 |  | Y21-S428 | 85 |  | Low |  |
| 24 | Smp_075280 |  | V19-T222 | 77 | LAMP-like protein | High |  |
| 25 | Smp_129840 |  | G25-S1161 | 206 |  | Medium |  |
| 26 | Smp_133270 |  | E34-T759 | 120 | Sel1-like protein | - |  |
| 27 | Smp_145420 |  | H30-I1733 | 282 | Plexin A3 | - |  |
| 28 | Smp_149740 |  | F26-T560 | 130 | Alzheimer disease beta-amyloid related | Medium |  |
| 29 | Smp_155810 |  | L29-T1100 | 184 | Protocadherin 11 | Medium |  |
| 30 | Smp_162520 |  | N20-P1070 | 162 | Protocadherin fat4 | Low |  |
| 31 | Smp_164760 |  | S22-P1160 | 184 | IgSF | Low |  |
| 32 | Smp_166300 |  | N27-I356 | 94 |  | High |  |
| 33 | Smp_168240 |  | Q25-T263 | 68 |  | High |  |
| 34 | Smp_171460 |  | L19-S841 | 161 | IgSF | Medium |  |
| 35 | Smp_176540 |  | V31-T956 | 146 | Protocadherin 18 | Medium |  |
| 36 | Smp_157070 |  | I19-P397 | 83 | EGF-domain-containing protein | Low |  |
| 37 | Smp_165440 |  | Q25-P363 | 74 | Netrin receptor unc5 | Low |  |
| 38 | Smp_136690 |  | N26-T674 | 118 | Acetylcholinesterase | Medium | [17] |
| 39 | Smp_061970 |  | F21-P518 | 95 | GPI ethanolamine phosphate transferase 2 | - |  |
| 40 | Smp_153390 | SmNNP5 | S19-S428 | 95 |  | Medium | [14] |
| 41 | Smp_072190 | SmLy6D (Sm29) | V27-T168 | 57 |  | High | [14] |
| 42 | Smp_064430 |  | A22-S155 | 53 |  | Medium | [19] |
| **secreted adhesion/growth factor/metabolite binding** |  |  |  |  |  |  |  |
| 43 | Smp_194840 |  | E18-I146 | 47 | NPC-like cholesterol-binding protein | High | [21] |
| 44 | Smp_194910 |  | N22-I180 | 51 | Saposin B domain-containing protein | Medium | [20] |
| 45 | Smp_063530 |  | E26-R193 | 49 | Apoferitin | Medium | [21] |
| 46 | Smp_141680 |  | I33-A659 | 110 | Fasciclin domain-containing protein | High | [14] |
| 47 | Smp_043650 |  | Q23-L81 | 37 | Prohormone npp-28 | Low |  |
| 48 | Smp_170550 |  | N27-Y928 | 164 | Granulin | Medium |  |
| 49 | Smp_035040 |  | N24-L241 | 63 | IgSF | Low |  |
| 50 | Smp_052660 |  | I22-R488 | 97 | Matabotropic glutamate receptor | - |  |
| 51 | Smp_128590 |  | K21-S260 | 69 | Laminin gamma3 | Low |  |
| 52 | Smp_135210 |  | I21-Q1584 | 220 | EGF-domain-containing protein | Low |  |
| 53 | Smp_136320 |  | H21-K154 | 48 | GSK3beta-interacting protein | Low |  |
| 54 | Smp_144130 |  | L26-V553 | 118 | Septate junction protein | Low |  |
| 55 | Smp_154760 |  | Q30-M2155 | 298 | EGF-domain-containing protein | Low |  |
| 56 | Smp_171780 |  | Q19-K260 | 61 | SPARC | High |  |
| 57 | Smp_178740 |  | Q31-V533 | 97 | Tesmin-related protein | Low |  |
| 58 | Smp_180600 |  | S22-S292 | 72 | IGF-binding protein | High |  |
| 59 | Smp_181220 |  | N18-I173 | 50 | C1q-binding protein | Medium |  |
| 60 | Smp_211020 |  | L32-Y703 | 122 | Discoidin domain-containing protein | - |  |
| 61 | Smp_016490 |  | E21-Q194 | 53 | SaposinB domain-containing protein | Medium |  |
| 62 | Smp_130100 |  | F20-I128 | 45 | Saposin domain-containing protein | Low | [20] |
| 63 | Smp_105420 |  | I19-S196 | 53 | Saposin domain-containing protein | High |  |
| 64 | Smp_105450 |  | Y19-C126 | 58 | Saposin domain-containing protein | Medium | [21] |
| 65 | Smp_202610 |  | V19-T135 | 46 | Saposin domain-containing protein | Medium |  |
|  | **Secreted protease / protease inhibitor** |  |  |  |  |  |  |
| 66 | Smp_090100 |  | E24-Q582 | 98 | Invadolysin | Low | [16] |
| 67 | Smp_067060 |  | H18-N340 | 73 | Cathepsin B1, isotype2 | High | [21] |
| 68 | Smp_103610 | SmCB1 (Sm31) | H18-N340 | 73 | Cathepsin B1, isotype1 | Low | [21] |
| 69 | Smp_019030 |  | D21-L455 | 91 | Cathepsin C/Dipeptylpeptidase 1 | Low | [21] |
| 70 | Smp_002600 |  | L17-G498 | 111 | Lysosomal Pro X carboxypeptidase | Medium | [21] |
| 71 | Smp_071610 |  | I24-L472 | 99 | Dipeptidyl-peptidase II | Medium | [21] |
| 72 | Smp_089670 |  | S19-N2127 | 409 | Alpha-2 macroglobulin | Medium | [15] |
| 73 | Smp_112090 | SmCE2a.3 | F19-I263 | 56 | Cercarial elastase 2a | - | [16] |
| 74 | Smp_119130 | SmCE1a.2 | W25-I263 | 55 | Cercarial elastase 1a | - | [16] |
| 75 | Smp_002150 | SmSP2 | S25-F501 | 97 | Trypsin-like serine protease | Low |  |
| 76 | Smp_141610 | SmCB2 | N23-N347 | 72 | Cathepsin B | Medium |  |
| 77 | Smp_147730 | SmKI-1 | Y21-K146 | 47 | Kunitz-type protease inhibitor | Medium |  |
| 78 | Smp_034420 | Sm12.8 | N20-S148 | 56 | Cystatin | Medium |  |
| 79 | Smp_075800 |  | Q20-G429 | 83 | Hemoglobinase | Low |  |
| 80 | Smp_210500 | SmCL3 | D17-V370 | 75 | Cathepsin L3 | Medium |  |
| 81 | Smp_132480 |  | E16-S393 | 79 | Subfamily A1A unassigned peptidase | Low |  |
| 82 | Smp_166280 |  | C20-L337 | 79 | Glutaminy cyclase | Medium |  |
| 83 | Smp_187140 |  | K21-F342 | 74 | Cathepsin L | Low |  |
| 84 | Smp_006510 | SmCE2a.2 | W25-I263 | 53 | Cercarial elastase 2a | - | [19] |
| 85 | Smp_090110 |  | E24-I591 | 103 | Invadolysin | - | [19] |
| **Secreted enzyme (not protease)** |  |  |  |  |  |  |  |
| 86 | Smp_145920 |  | Y30-F340 | 71 | Protein tyrosine sulfotransferase | Low |  |
| 87 | Smp_040790 |  | E24-E213 | 54 | Peptidyl prolyl cis-trans isomerase B | Medium |  |
| 88 | Smp_008320 |  | D18-D325 | 72 | Pap-inositol-1,4-phosphatase | Low |  |
| 89 | Smp_021730 |  | G30-D220 | 52 | Cytochrome c oxydase subunit Vb | Low |  |
| 90 | Smp_078800 |  | E25-S192 | 49 | DnaJ subfamily B | Low |  |
| 91 | Smp_007450 |  | R22-G160 | 48 | Heat-shock 67b2 | Medium |  |
| 92 | Smp_026930 |  | T51-V442 | 85 | Acetylglucosaminyltransferase | Medium |  |
| 93 | Smp_059910 |  | D28-Q274 | 70 | Ser-Thr protein phosphatase | - |  |
| 94 | Smp_089240 |  | K43-Y498 | 93 | Acetylglucosaminyltransferase | - |  |
| 95 | Smp_134800 |  | K20-T1590 | 242 | Tyr protein kinase | Low |  |
| 96 | Smp_018760 |  | N25-L991 | 152 | Alpha glucosidase | Low | [19] |
| **Secreted venom/secreted MEGs/short** |  |  |  |  |  |  |  |
| 97 | Smp_194860 | Sm8.7 | E20-E92 | 39 |  | Medium | [17] |
| 98 | Smp_138080 | MEG-3.1 | A21-S146 | 50 |  | High | [17] |
| 99 | Smp_194830 | SmKK7 | K21-D79 | 37 |  | High | [16] |
| 100 | Smp_001890 | VAL18 | K27-Y194 | 55 |  | Low | [16] |
| 101 | Smp_002070 | VAL4 | K22-E181 | 57 |  | Low | [16] |
| 102 | Smp_138060 | MEG-3.3 | A21-G151 | 46 |  | High |  |
| 103 | Smp_180620 | MEG-17 | N17-R65 | 41 |  | High |  |
| 104 | Smp_123540 | VAL12 | I22-L24 | 57 |  | Low |  |
| 105 | Smp_123550 | VAL8 | Q24-K261 | 66 |  | Medium |  |
| **Secreted - no known function** |  |  |  |  |  |  |  |
| 106 | Smp_181070 |  | K25-Q114 | 43 |  | Low |  |
| 107 | Smp_004710 |  | M26-S127 | 42 |  | High |  |
| 108 | Smp_061310 |  | Y20-E140 | 65 |  | Low |  |
| 109 | Smp_005060 |  | E17-I146 | 45 |  | - |  |
| 110 | Smp_141500 |  | N27-L121 | 56 |  | High |  |
| 111 | Smp_006060 |  | P35-P380 | 92 |  | High |  |
| 112 | Smp_063330 |  | K24-F182 | 48 |  | High |  |
| 113 | Smp_096790 |  | T21-E94 | 38 |  | Medium |  |
| 114 | Smp_201730 |  | E24-R92 | 38 |  | Medium |  |
| 115 | Smp_019000 |  | D28-D236 | 53 |  | Medium | [19] |

### Recombinant protein expression using a mammalian expression system

For membrane anchored proteins, the entire ectodomain was selected and the signal peptide removed using predictions from SignalP v3.0 [34]. Corresponding cDNAs were made by gene synthesis and codon-optimised for expression in human cells (Invitrogen, GeneArt). The ectodomains were flanked by unique *Not*I and *Asc*I restriction enzyme sites and subcloned into an expression plasmid containing a high-scoring signal peptide [29], and a C-terminal tag containing rat Cd4d3+4 domain, a BirA monobiotinylation sequence, and 6-His tag (Addgene plasmid# 50803) [36]. All expression constructs have been deposited with Addgene (Addgene plasmids #120590 to 120704). Plasmids were transiently transfected in HEK293-E or HEK293-6E cells [37] and supernatants collected as previously described [38], and detected by Western blotting with 0.02 µg/mL streptavidin-HRP (Jackson Immunoresearch) as described [29]. Where proteins showed signs of proteolytic cleavages, transfections were repeated in the presence of a protease inhibitor cocktail (Sigma).

### Human sera from endemic regions infections

A pool of sera from ten Ugandan adults living in a high-transmission area showing high soluble worm antigen IgG1 responses and treated with praziquantel [39, 40] or non-exposed European control sera were used to initially characterise the proteins. All patient samples were collected in accordance with the Uganda National Council for Science and Technology (UNCST) and the Cambridge Local Research Ethics Committee.

### Sera from experimental mouse infections

The life cycle of the NMRI (Puerto Rican) strain of *Schistosoma mansoni* was maintained by routine infections of mice and susceptible *Biomphalaria glabrata* snails under the UK Home Office Project Licences Nos. P77E8A062 and PD3DA8D1F, and all protocols were approved by the local Animal Welfare and Ethical Review Body (AWERB). Seven-week-old BALB/C female mice were infected percutaneously by tail immersion in water containing 200 cercariae for 40 minutes under general anaesthesia, or by injection of 350 cercariae intraperitoneally. Blood samples were collected at 8, 21 and 42 days post-infection.

### Enzyme-linked immunosorbent assays (ELISAs)

Protein expression was quantified by ELISA as described [29]. Briefly, biotinylated proteins were diluted in HBST/2%BSA and captured on streptavidin-coated microtitre plates and detected by mouse anti-rat Cd4 OX68 antibody (AbD Serotec), followed by an alkaline-phosphatase-conjugated anti-mouse secondary antibody (Sigma). Based on the ELISA readings, proteins were classified into high (typically >5 µg/mL transfection supernatant), medium (between 1 and 5 µg/mL transfection supernatant) and low (<1 µg/mL transfection supernatant) levels of expression. To determine the presence of heat labile epitopes, biotinylated proteins were captured on streptavidin-coated plates either untreated or following heat-treatment for 10 minutes at 80°C before being incubated with sera at 1:1,000 dilution in HBST/2%BSA. Sera from experimentally-infected mice were diluted 1:1000 in HBST/2%BSA and binding was detected with a horseradish peroxidase (HRP)-conjugated anti-mouse secondary antibody recognising IgA, IgM and IgG. Analysis was performed for each individual by reference-subtracting the OD value of the pre-infection samples from the OD readings at all subsequent time points. OD values for each protein were compared to that of a negative control and seropositivity was defined as OD_protein_ > OD_control_ + 3SD_control_.

## Results

### Selection and expression of a panel of 115 secreted and cell-surface proteins from Schistosoma mansoni

To compile a library of recombinant *S. mansoni* proteins that could be used for immunodiagnostics screening, we first made use of published proteomics data from the cercarial, schistosomula or adult stages. We identified those with features of cell surface and secreted proteins and selected a panel of 36 proteins (Table 1). Because of the difficulty in solubilising membrane-embedded proteins and the fact that glycosylated peptides are often uninformative in mass-spectrometry-based approaches, we sought to further enrich our library using transcriptomics data. To identify genes transcribed early after infection, we selected 1,302 transcripts enriched at 48 versus 3 hours post-infection, and the 1,000 most abundant transcripts in the 48-hour schistosomula. Proteins likely to be secreted or membrane-anchored were identified from these sets based on the presence of a N-terminal signal peptide or membrane-anchoring sequences, resulting in a list of 274 genes. Since the correct folding of recombinant proteins critically depends on correct gene structure, manual curation was performed on all candidates to check for completeness of their open reading frames. After further removal of mitochondrial, endoplasmic reticulum and multiple-pass membrane proteins, 79 secreted or membrane-anchored candidates with correct gene structures were added to the shortlist, resulting in a final set of 115 genes (Table 1). Approximately one third of the selected genes encoded single-pass transmembrane or GPI-anchored proteins, while the remaining two thirds were secreted proteins and enzymes. Based on sequence similarities with other proteins, these secreted proteins and enzymes were further divided into five broad categories: secreted adhesion and metabolite-binding proteins, secreted proteases, other secreted enzymes, secreted venom allergen-like (VAL) and micro-exon genes (MEG), and finally putative secreted proteins with no known homology. Proteins were expressed in HEK293 cells, quantified by ELISA, and their integrity determined by Western blotting (Figure 1). The majority of the proteins produced could be detected at their expected full-length size including the very large proteins Sm200 (257kDa, Protein 4) and α2-macroglobulin (409kDa, Protein 72), and only twelve proteins (10%) could not be detected by ELISA or Western blot (Table 1, Figure 1). Using a mammalian expression system, we were therefore able to successfully produce over a hundred recombinant *S. mansoni* surface and secreted proteins, which will be a valuable resource for functional and immunological studies.

**Figure 1.**
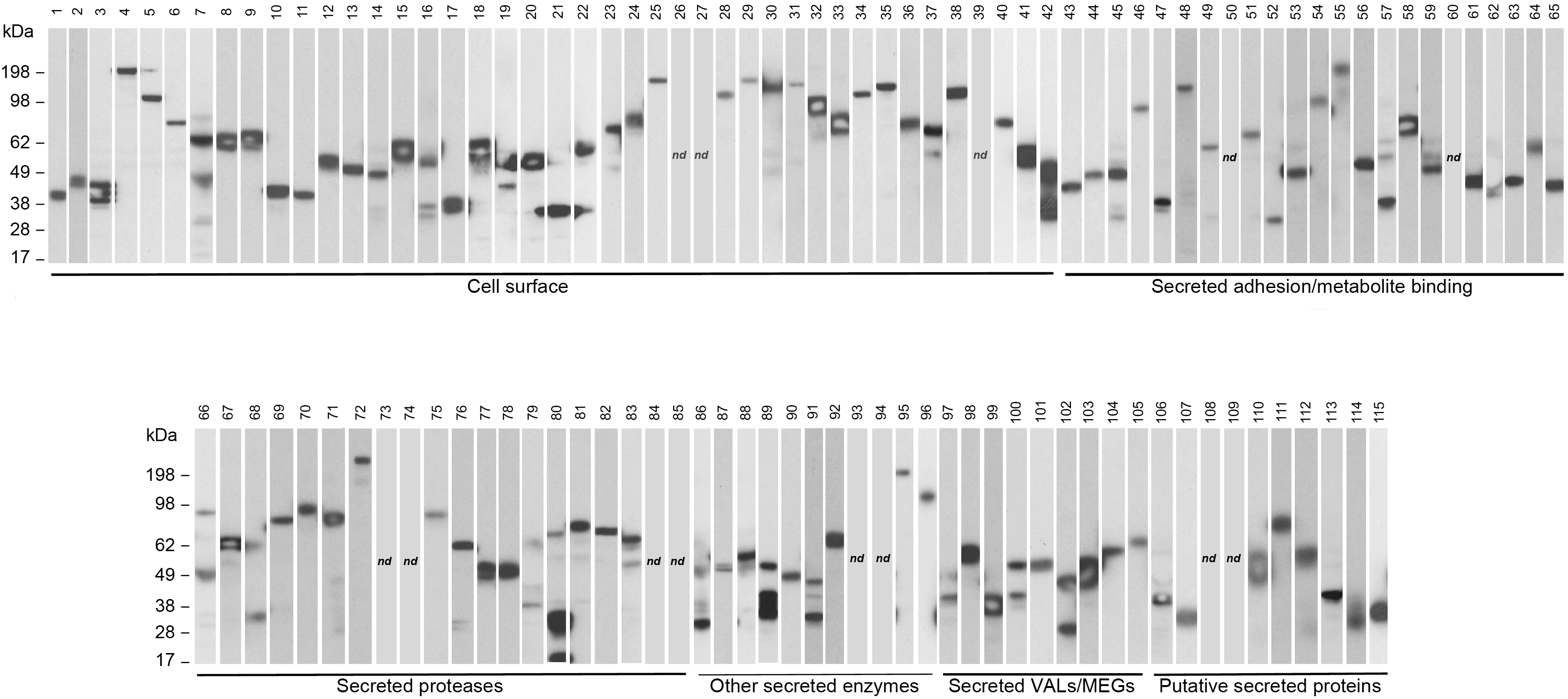
Expression of a library of 115 recombinant cell-surface and secreted proteins from *Schistosoma mansoni* expressed as secreted enzymatically monobiotinylated recombinant proteins in HEK293 cells. Supernatnants were resolved by SDS-PAGE under reducing conditions and blotted. Proteins were detected with streptavidin conjugated to horseradish peroxidase. Around a third of the protein library consists of membrane-tethered surface proteins, while the remainder of the library corresponds to secreted proteins as indicated. The expected molecular masses are indicated in kilodaltons (kDa) on the left-hand side of each panel. nd, not detected by Western blot.

### S. mansoni recombinant proteins expressed in mammalian cells contain heat-labile conformational epitopes

Many antibodies that are elicited in the context of a natural infection recognise conformational epitopes displayed on the native protein. Since most of the proteins selected in the library have no known function or binding partners, we determined which fraction of the proteins contained conformational epitopes by comparing the immunoreactivity of pooled sera from ten adult Ugandan patients living in a high-transmission area to untreated and heat-treated proteins (Figure 2). All proteins except numbers 18 and 57 were seropositive, and sixty-six of the expressed proteins (64%) were highly immunoreactive with an OD reading > 0.3 (Figure 2). The majority of proteins showed moderate to strong loss of immunoreactivity after heat-treatment with only twelve proteins showing little or no loss of reactivity, suggesting that these latter proteins were either natively unstructured, misfolded, or contained heat-stable domains (Figure 2). In summary, the majority of the recombinant proteins produced were reactive against immunoglobulins from individuals living in high-endemicity areas, who have experienced infections with *S. mansoni*. The decrease in reactivity observed after heat-treatment suggest that most of these recombinant proteins contain conformational epitopes that are also present in native *S. mansoni* proteins and are recognised by host antibodies in the context of a natural infection.

**Figure 2.**
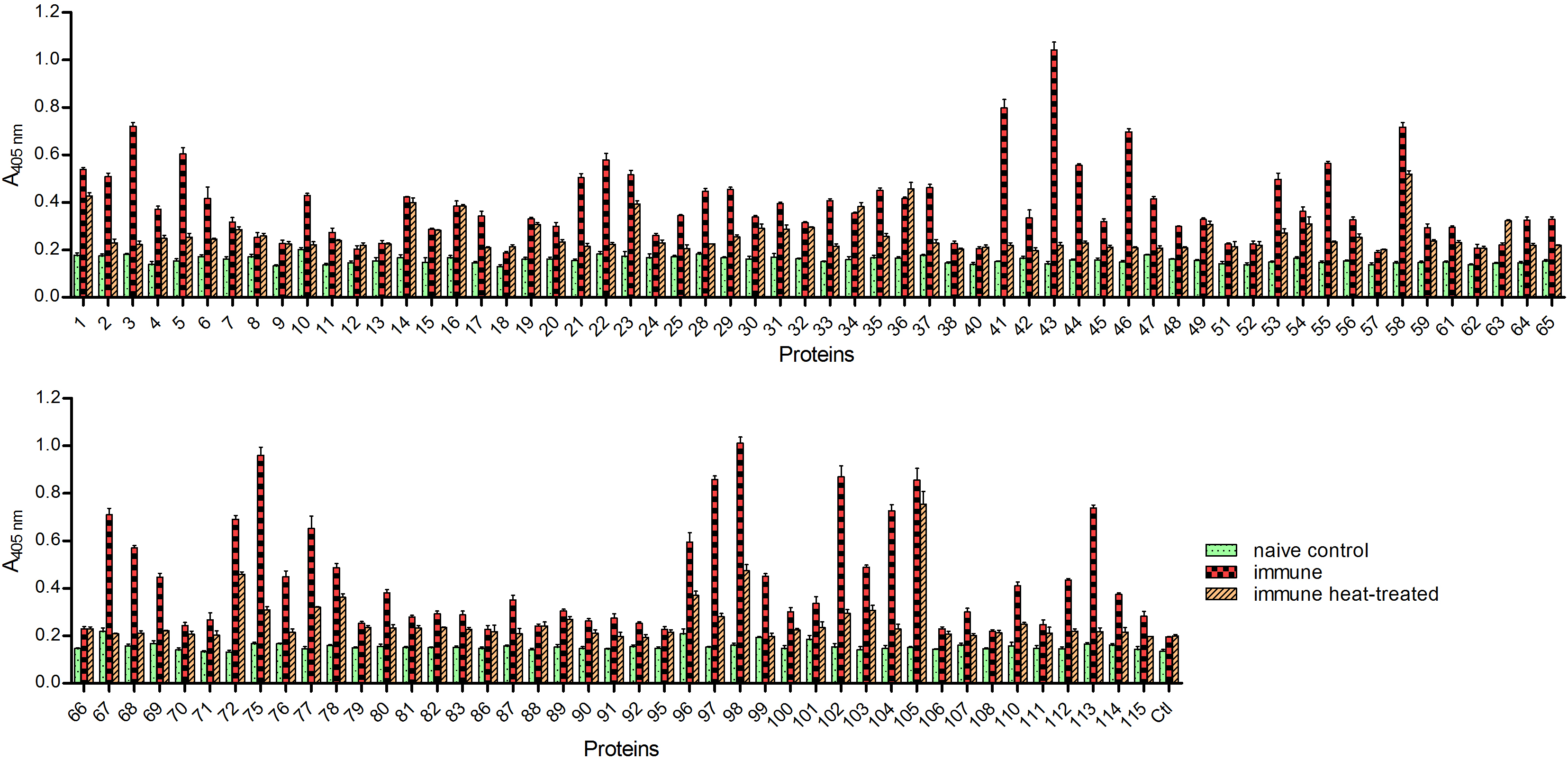
The majority of recombinant proteins are immunoreactive to sera from individuals living in schistosomiasis-endemic areas and contain heat-labile conformational epitopes. Recombinant proteins were probed with pooled sera from individuals living in schistosomiasis-endemic areas (“immune”, red checkered bars) or individuals from the UK who have never been infected (“naïve”, green dotted bars). To test for the presence of heat-labile epitopes, recombinant proteins were also heat-treated to 80°C for 10 minutes before being exposed to immune sera (“immune heat treated”, orange hatched bars). Most proteins were immunoreactive when exposed to sera from endemic areas and contained heat-labile epitopes as shown by a decrease of reactivity following heat-treatment. All measurements were performed in triplicate; error bars = SD.

### The humoral response elicited by mixed infections in mice identifies cathepsins and saposin-domain-containing proteins as early markers of infection

Sera from adults living in high-transmission areas and who have received praziquantel treatment are expected to have a broad repertoire of antibodies against the *S. mansoni* parasite. Although these sera were useful in characterising our recombinant protein library, they would not be suitable to provide any information about the kinetics of the host immunological response in the context of a primary infection. Identifying antigens that are first recognised by the host immune system could be particularly useful for the identification of new early markers of exposure to the parasite. To address this in a controlled experimental setting, we used mice as an animal model of infection. Sera were collected at 8, 21 and 42 days post-infection from inbred female BALB/C mice infected either percutaneously with 200 cercariae or through intraperitoneal injection of 350 cercariae. Sera from three individual mice infected percutaneously or a pool of sera from three mice infected through the intra-peritoneal route were analysed and compared (Figure 3). None of the antigens showed consistent reactivity across samples at 8 days post infection, as might be expected from the initial exposure to a pathogen. However, reactivity to antigen 44 (and to a lesser extent, antigen 3) could be detected across all samples from 21 days post-infection. At 42 days, a further eight antigens showed reactivity across all sera tested (Proteins 10, 41, 62, 63, 64, 67, 68, and 106) and all but one mouse showed strong reactivity to protein 71. Overall, reactivity with the pool of sera from mice infected intraperitoneally was stronger than individual mice infected percutaneously, which might be a reflection of the higher number of cercariae used. Proteins 44, 62, 63, 67 and 68, which all belong to either the saposin-containing or cathepsin families, were the most immunoreactive, along with protein 106. In addition, reactivity to three Ly6 family members Ly6F, Ly6B and Ly6D (proteins 3, 10, and 41, respectively) was also detected. Overall, however, members of the saposin-containing and cathepsin families appeared to be the most immunoreactive proteins in the murine model at a relatively early stage of infection.

**Figure 3.**
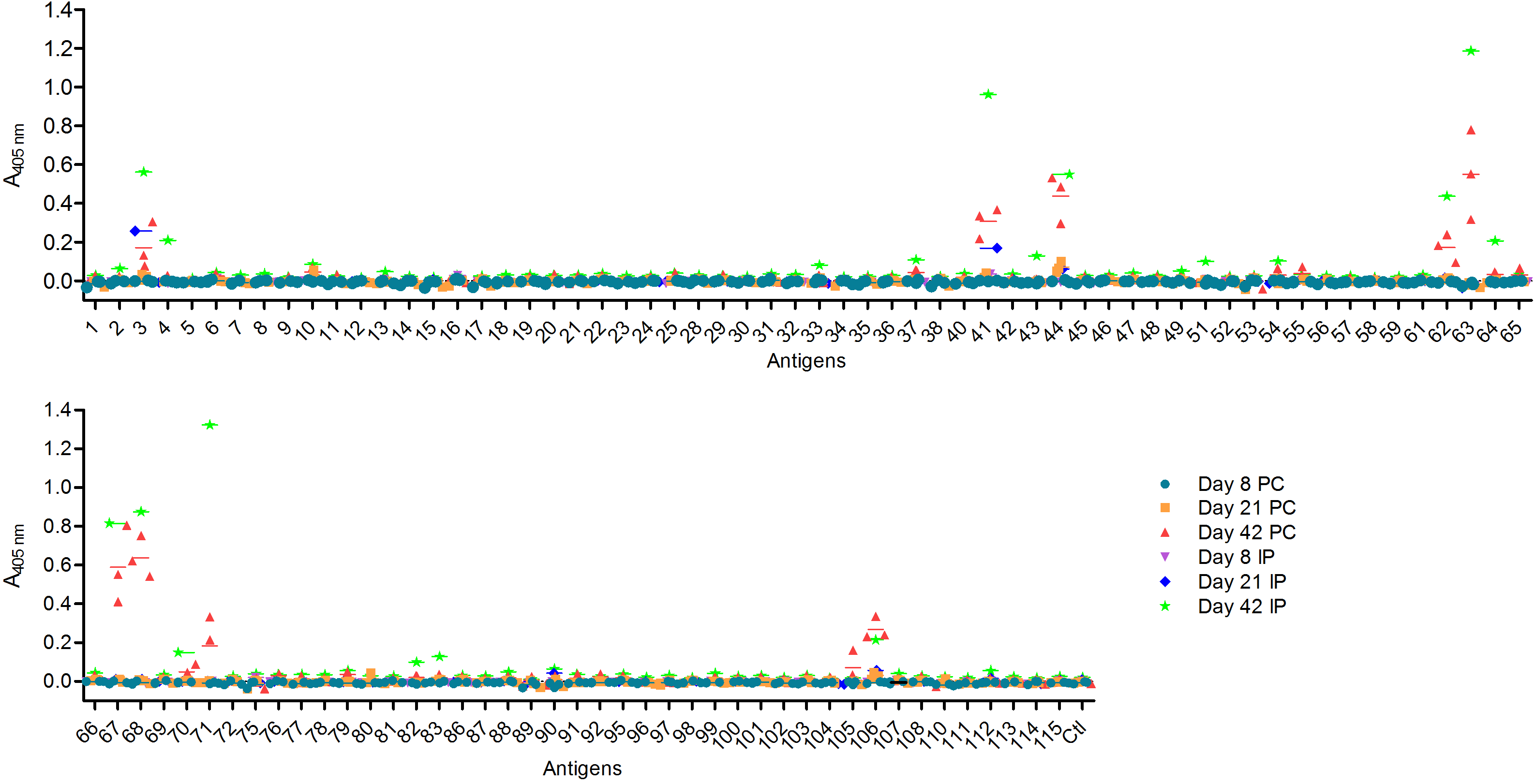
Analysis of the acquired antibody response to *S. mansoni* antigens in experimentally controlled infected mice. Individual sera from three mice infected percutaneously or pooled sera from three mice infected intraperitoneally were analysed at 8, 21 and 42 days post-infection. The aim was to try and capture the antibody reactivity at different stages of parasite maturation: schistosomule at 8 days, immature adult at 21 days, and mature adult at 42 days. Each data point represents the average of triplicate experiments; lines show the mean of the data for animals infected either percutaneously or intraperitoneally.

## Discussion

Despite its widespread distribution and high morbidity rate, schistosomiasis remains a neglected tropical disease whose true incidence and health impact likely remains underestimated. Here, we have described a recombinant protein library containing 115 secreted and cell-surface proteins from different developmental stages of *Schistosoma mansoni*. We used this new valuable resource to determine the kinetics of the mouse humoral response to *S. mansoni* infections. While other large arrays of parasite proteins have been used for diagnostic and immuno-epidemiological purposes, they have mostly relied on cell-free or bacterial expression systems [41–43]. Mammalian expression systems are more suitable for the addition of structurally important posttranslational modifications found on extracellular proteins and thereby preserve conformational epitopes that can be recognised by antibodies. Using this approach, we successfully expressed over one hundred of these proteins, the majority of which were observed at their expected size. Although the twelve proteins that could not be expressed were equally distributed amongst the different protein families represented in the library, three of them corresponded to elastases. Using sera from patients living in a high-transmission region of Uganda, we observed that 64% of the expressed proteins were considered to be strongly immunoreactive (OD>0.3) with the majority (82%) showing sensitivity to heat-treatment, an indicator of tertiary folding. Although the lower immunoreactivity for 37 proteins may suggest incorrect folding, they could also be weakly immunogenic in humans, or not directly exposed to the host immune system. Although a large fraction of the natural host immune response has been reported to be directed against parasite glycans present at different stages of development [44–46], the conformation-sensitive immunoreactivity that we observed in our assay is almost certainly directed against the protein backbones since the immunoreactivity against glycans would not be affected by heat-treatment and the glycan groups added on the recombinant proteins in HEK293 cells may differ substantially from the ones present on endogenous proteins. Of the sixteen most immunoreactive proteins observed, eleven had already been identified in the surface and secreted proteome of *S. mansoni*; by contrast, only six out of the 37 proteins with low serum reactivity (16%) have already been described in proteomics studies. The use of a single mammalian expression system for the production of large panels of recombinant proteins is particularly attractive for the systematic comparison of antigens in diagnostic, immuno-epidemiological or vaccination studies and we have used this approach previously for the identification of potentially protective antibodies against malaria [29, 31, 36, 47].

Mass administration of praziquantel to schoolchildren has been the mainstay of control programmes against schistosomiasis and this has proved relatively successful in reducing parasite burdens and contributing to elimination from some areas. Continued surveillance and early detection of new cases remains critical to avoid any risk of resurgence; however, the commonly used Kato-Katz method is not sensitive enough to detect low levels of infection, resulting in the underestimation of the number of cases [9]. Detection of parasite-derived glycans such as the circulating anodic antigen (CAA) in the urine or serum of patients by lateral-flow test is currently considered the most sensitive point-of-care assay for the detection of current *Schistosoma* infections as reactivity disappears rapidly following praziquantel treatment. In areas where schistosomiasis has been eliminated or is close to elimination, alternative methods of surveillance might be needed. By persisting several months or even years after infection [48], antibody responses provide a useful historical measure of parasite exposure and are better suited for monitoring populations at risk of resurgence. Currently, most host antibody responses are measured against crude parasite preparations such SEA or SWAP, which suffer from considerable cross-reactivity with other helminths antigens. The use of species-specific recombinant proteins as diagnostics could therefore be more reliable.

A striking feature of this study is the relatively small number of antigens eliciting a patent immune response in the few weeks following a primary infection, which could reflect the ability of the parasite to evade the host immune response. Strikingly, the antibody responses observed at 6 weeks were dominated by saposin-containing and cathepsin proteins: out of the ten proteins that showed reactivity in all animals at 6 weeks post-infection, four were saposin-domain-containing proteins and two were cathepsins. Proteins 66 and 67 are two isoforms of the Cathepsin B1 protease that has been suggested to have inbuilt adjuvanticity [49]. Interestingly, SjSAPLP1 and SjSAPLP4, the *Schistosoma japonicum* orthologs of proteins 44 and 63 respectively, have been proposed as markers of infections in mice and humans [50, 51], and it is therefore likely that *S. mansoni* orthologs of saposin-domain-containing proteins will be early markers of infection in human sera too. Both saposins and cathepsins are produced in the parasite’s gut and have been involved in the digestion of lipids and proteins [52, 53] but their transcriptional up-regulation at the cercarial and day3 schistosomula stages suggest they might also play a role in the gut of the parasite larvae [26]. Their exposure to the humoral system of infected individuals suggests they are regurgitated by the worm during the feeding process. While several saposin-domain-containing proteins present in the library were immunoreactive, their sequence identity ranged between 9 and 28%, making cross-reactivity unlikely.

Another group of immunoreactive proteins consistently observed across samples were members of the uPAR/Ly6 domain-containing family, whose expression has been reported at the schistosomule and adult stages [54–56]. In our study, reactivity to three members (Proteins 3, 10 and 41) corresponding to Ly6F, B and D, respectively was observed in mouse serum samples. These proteins have previously been shown to elicit strong antibody responses in rat, mouse and human sera [56]. Members of this family share some homology with the complement-inactivating Cd59 protein although they do not seem to have preserved this function [54].

Quantitation of gene expression suggests that these genes are, however, already actively transcribed at the schistosomula stage [15, 54, 55], several weeks before reactivity from host antibodies can be detected. The ability of schistosomes to manipulate and evade the host immune system has been described although the specific proteins and mechanisms orchestrating this escape are not yet clearly defined [57–60].

By producing our recombinant proteins in a mammalian expression system, we have paid particular attention to their correct folding, and thus this new resource could be used in a wide range of cellular and molecular assays such as vaccine screening, cellular assays looking at immunomodulatory functions, immunoepidemiological studies, or the identification of host binding partners by receptor-ligand screening. We envisage these proteins will be useful to the wider scientific community to further understand *Schistosoma* biology.

## Acknowledgements

We would like to thank Catherine McCarthy for technical help with the mouse infections and David Dunne for helpful discussions and comments on the design of the study. This work was supported by the Wellcome Trust grant 206194.

